# Antimicrobial activity of *Syzygium aromaticum* (clove) against *Staphylococcus aureus* and *Listeria monocytogenes* is enhanced by optimized extraction methods

**DOI:** 10.64898/2026.01.06.698043

**Authors:** C.H Justice-Alucho, W Braide

## Abstract

Clove (*Syzygium aromaticum*) is a food spice rich in phenolic profile with potent antimicrobial properties. However, the influence of solvent extraction efficiency and antimicrobial performance against selected foodborne pathogens needs to be further elucidation. In this study, four solvents (80% ethanol, 80% acetone, 75% methanol, and water) were analyzed for their impact on clove bud extraction yield and antimicrobial activity against *Staphylococcus aureus* and *Listeria monocytogenes*. Extraction yield varied significantly across solvents, with 80% acetone producing the highest recovery, while water yielded the lowest, with no significant difference from 75% methanol (p > 0.05). Preliminary agar diffusion screening revealed inhibition zones for all extracts and Minimum inhibitory concentration (MIC) and minimum bactericidal concentration (MBC) assays demonstrated that acetone and ethanol extracts exerted the strongest inhibitory effect on *S. aureus*, whereas ethanol and methanol extracts were most effective against *L. monocytogenes*. Water extract consistently exhibited the weakest antimicrobial activity. A perfect correlation (r = 1.00) was observed between MIC and MBC for both organisms, and a strong negative correlation existed between extraction yield and antimicrobial thresholds for *S. aureus* (r = −0.98; p < 0.05), suggesting solvent-driven phytochemical concentration as a determinant of bioactivity. Collectively, these findings reaffirm the critical role of solvent polarity in maximizing phenolic extraction and antimicrobial potency of clove extracts as a promising natural antimicrobial agent for the control of foodborne pathogens.

## Introduction

The demand for natural and safer alternative therapies in the treatment of disease has been on the rise in recent times. Bioactive compounds have been shown to exert health promoting properties like anticancer, antidiabetic, anti-inflammatory, antibacterial antiviral and prebiotic features (Uwineza & Waśkiewicz, 2020). These bioactive compounds have found several applications in the cosmetic, food, feed, pharmaceutical and agricultural industries(Abdul Aziz et al., 2023). Bioactive compounds from clove (*Syzygium aromaticum)* have been proven by studies to exert antimicrobial and antioxidant activities making it an important nutraceutical and functional food (Abdul Aziz et al., 2023). Although clove is a traditional food spies recognized across Asia and Africa, it is widely used as a nutraceutical given its antioxidant, and antimicrobial activities. Several studies have emphasized its antifungal, antiviral, anticarcinogenic and preservative properties (Cortés-Rojas et al., 2014). These properties are said to conferred by eugenol which is the major bioactive compound ranging from 9381.70 to 14650.00 mg per 100 g. It also possesses flavonoids, phenolic acids, gallic acid, kaempferol, quercetin essential oils, hydroxybenzoic acids, hydroxycinnamic acid and hydroxyphenyl propene (Neveu et al., 2010). Furthermore, clove also contains polyphenolic components which also confer antioxidant properties making it an important potential pharmaceutical component (Pérez-Jiménez et al., 2010). Its antioxidant activity makes it an important food preservative as ethanol and aqueous extracts of clove exerts superoxide radical capture and scavenging of DPPH potential (Dudonné et al., 2009). 1% of clove extract have been shown to exhibit inhibitory activity against fungi like *Candida albicans, Mucor* sp, *Microsporum canis, Trichophyton mentagrophytes* and bacteria including *Escherichia coli, Staphylococcus aureus, Bacillus, Pseudomonas aeruginosa* and *Proteus vulgaris*. Moreover, clove has been shown to inhibit the growth of 31 strains of *Helicobacter pylori* after 9-12 hours of incubation even at very low pH (Ali et al., 2005). Clove analgesic ability is conferred by the eugenol through the activation of calcium and sodium channels in ganglionic cells and receptors (Li et al., 2008). Clove’s antiviral activity against Ebola, herpes simplex virus type 1 and 2, and influenza virus (Lane et al., 2019) (Dai et al., 2013). The eugenol derivative may also have inhibitory effect on West Nile, zika virus, dengue virus, yellow fever and HIV-1 by limiting viral replication via increased lymphocyte proliferation (Behbahani et al., 2013). (Haro-González et al., 2021). Clove has also been utilized in recent times as sources of biofuels and a good source of biodegradable plastic further emphasizing its potential application in environmental health (Varghese et al., 2023)

Different extraction methods are available for bioactive components of clove buds including solvent extraction, Soxhlet, maceration, reflux extraction and distillation. (Abdul Aziz et al., 2023). The yield and purity of the bioactive compounds is affected by these different methods and the extraction solvent. The dielectric constant of liquid solvents are decreased when the temperature and pressure is high leading to an increase in the rapid solubility and extraction of target compounds in the matrix into the extracting solvent (Santana et al., 2023).

*Staphylococcal* infections have evolved into several life-threatening infections including sepsis, pneumonia, endocarditis, bronchitis, impetigo and enterotoxin intoxication (Vestergaard et al., 2019). These infections are treated using antibiotics that target the cell wall synthesis, translation, transcription, and DNA synthesis in the bacteria (Vestergaard et al., 2019). With the rise in global antimicrobial resistance resulting from excessive abuse of antibiotics, *Staphylococcal* infection treatment has been threatened calling for the urgent need for alternative and sustainable therapies to prevent and treat these infections. The emergence of Methicillin Resistant Staphylococcus aureus (MRSA) strains encoded by the *mecA* and *mecC* genes as well as penicillin resistant *S. aureus* encoded by the *blaz* gene, which have been shown to be resistant to beta-lactams up to 1600ug/ml (Parvez et al., 2008). These realities call for an urgent need for a sustainable and more efficient therapy to curb antibiotic resistance infections.

*Listeria monocytogenes* mediated infections are transmitted to humans via food infection such as dairy products. It is the causative agent for gastroenteritis with a 20-30% mortality rate (De Noordhout et al., 2014). Listeriosis infection is prevalent in immunocompromised patients and pregnant women leading to spontaneous still birth, preterm birth or abortions(De Noordhout et al., 2014) While sometimes proliferate in pasteurized final food products causing an outbreak even after adequate control have been put in place, controlling preservation conditions of the final food product might provide increased shelf-life without contamination even at refrigeration temperatures (Cava et al., 2007a). Listeria outbreak have been recorded in the united states in cantaloupe Colorado in 2100 killing 33 people while infecting 147 people (Gillespie et al., 2010). Very recently, industries are incorporating natural products as antimicrobial agents to improve aroma, flavor and shelf life of food products (Cava et al., 2007b). Listeria ability to survive different mammalian host is conferred by its adaptation mechanisms. It colonizes the intestinal epithelium when ingested by the host, and move to the lamina propria and then disseminates through the lymph node and blood to the organs (Radoshevich & Cossart, 2018). In immune cells, *L. monocytogenes* is internalized in phagocytic and non-phagocytic cells by the receptor mediated endocytosis mediated by internalin A and B, E-cadherin and Met, and hepatocyte growth factor (Radoshevich & Cossart, 2018) (Cossart & Helenius, 2014). Several studies have highlighted the effect of bioactive compounds from herbs on *Listeria* proliferation in food through targeted virulence approach and antimicrobial activities. The antimicrobial activities of different solvent extracts of clove on selected foodborne pathogens have not been fully elucidated hence this study aims to understudy the efficacy of extraction methods of clove buds in inhibiting the growth of *Staphylococcus aureus* and *Listeria monocytogenes*.

## Materials and Methods

### Collection of plant material

Clove buds were purchased from relief market in Owerri, Imo state Nigeria.

### Preparation of the solvent extracts of clove buds

Cloves were washed, oven dried and grounded into fine powder using a sterile coffee grinder after which 5g of the milled clove buds were weighed into 100 mL of 75% acetone, 80% methanol, 80% ethanol and water in a sterile conical flask. The metabolites extraction was done following methods by (Herald et al., 2012) with some minor modifications. Briefly, the mixture was shaken for 2 hours at room temperature at 120 rpm after which they were incubated at -20^0^ C for 24 hours. Subsequently, the mixture was centrifuged at 3000 x g for 10 minutes and the supernatant was collected using a sterile filter paper. The supernatant was added in pre-weighed sterile beakers and oven dried at 30^0^ C for less than 24 hours. Following drying, samples were weighed and reconstituted with 20% sterile DMSO using a volume appropriate to obtain a concentration of 200mg/ml of the extract. The percentage of the DMSO was determined after a prior testing on each isolate following recommendation from (Eloff, 2019). Samples were stored in -20^0^ C until they were further analyzed.

### Test isolate and Inoculation procedure

*Staphylococcus aureus* and *Listeria monocytogenes* laboratory strain was grown overnight in Tryptose Soy Broth (TSB) and Tryptose Soy Broth-Yeast extract (TSB-YE) respectively under shaking at 120 RMP at 37 ^0^ C for approximately 12 hours to achieve a logarithmic growth of the isolates (Eloff, 2019). Following incubation, the broth cultures were centrifuged at 5000 x g for 4 minutes to obtain bacterial pellets. The pellets were resuspended in 5 mL of 0.85% saline following two pellet wash steps. The bacterial densities were adjusted to on Optical Density of 0.5 ca equivalent to 10^8^ CFU/ mL. Two 10-fold serial dilutions of the suspension was done using sterile TSB broth after which it was incubated for two hours to enable bacterial reach logarithmic growth prior to analysis.

### Preparing dilutions of the extracts

Double fold dilutions of the extracts were sequentially made in 96-well plate by adding 100 uL of the extract to 100 uL of Muller-Hinton broth (MHB) in the first well to obtain a 2x dilution of the extract (Muraina et al., 2009). Next, 100 uL of the 2x extract dilution was added to another well containing 100 uL of MHB to obtain a 4x dilution and this was continued until a 64X dilution of the extract was obtained. Quadruplicates of each sample was made for all solvent extraction types (Muraina et al., 2009).

### Determination of Yield extract percentage for each extraction method

The percentage yield of the extract was obtained for each extractant by dividing the weight of the extract by the weight of the starting material and multiplying by 100 (Nur et al., 2020.).

### Antimicrobial screening by zone of inhibition

Following reports on the antimicrobial activities of clove ethanolic extracts on bacteria, we assessed the difference in antibacterial activity of clove following different extraction solvents. To do this we started by testing their activity via zone of inhibition method. This method was adopted from (Nwokafor et al., 2020) with some minor modifications. 100 uL of 1.0 × 10^6^ CFU/mL of each bacterial suspension was inoculated on Muller Hinton agar plates and with a sterile swab, the inoculum was evenly spread on the plate. Sterile blank discs of about 6mm-diameter were impregnated with 20 uL of each extract and placed at an equidistance on the seeded agar plate. The volume of the extract was specified following recommendations from (Eloff, 2019). This was done in three independent replicates with a DMSO impregnated disc as DMSO control and blank disc as negative control and 30ug ciprofloxacin and ampicillin antibiotic disc were used as antibiotic controls for *S. aureus* and *L. monocytogenes* respectively. The plates were incubated for 24 hours at 37^0^ C.

### Quantification of Antimicrobial activity

To quantify the antimicrobial activities of the extracts, minimum inhibitory concentrations (MIC) and Minimum Bactericidal Concentration (MBC) of the extracts on each isolate was assayed using microbroth dilution method as described by (Parvekar et al., 2020) with minor modifications. In a 96 well plate, 100 uL of the double fold diluted extracts was added into each well. Thereafter, 10 uL of 1.0 x 10^6^ CFU/mL of the isolates were inoculated into each well to obtain a final bacteria density of 1.0 x 10^5^ CFU/mL in each well. Sterility and positive control wells containing only the extract and media without the isolates and media and bacteria only were provided respectively. The plates were incubated at 37^0^ C for 18 – 24 hours after which 20 uL of 0.01% resazurin was added into each well and incubated in the dark for 2 hours to develop a color change. Wells which changed the color of the resazurin from blue to pink were considered negative while the least concentration of the extract that did not change from blur to pink was considered the MIC. Thereafter, 100 uL from each well was plated on Tryptose Soy Agar (TSA) plates and incubated using the same conditions. the least concentration of the well that did not produce any visible colonies on the plates was considered the MBC.

## Results

### Extractant have impact on extraction yield of clove bud extract

The extraction yield of clove bud from different extractants including 80% ethanol, 80% acetone, water and 75% methanol are shown in table 1. The maximum extraction yield was observed for 80% acetone, and the least yield was observed for water. Moreover, there was no significant difference seen between water and 75% methanol (p>0.05).

**Figure 1: Table 1:** percentage yield of sample extracts of clove buds from the different extracts.

| Extraction solvent | Weight of extract (g) | Extraction yield (%) |
| --- | --- | --- |
| 80% ethanol | 0.44±0.15 | 8.66±2.7 |
| 80% acetone | 0.47±0.02 | 9.32±0.5 |
| 75% methanol | 0.35±0.11 | 6.99±2.1 |
| Water | 0.27±0.01 | 5.33±0.1 |

The data represent mean ± standard deviations of three independent replicates. $p > 0.05$ for ethanol and acetone indicating there is no significant difference in extraction yields of both extraction methods. $P < 0.05$ for all extraction solvents and water indicating a significant difference.

### Qualitative screening of extracts on isolates

To identify if the clove bud extracts have any antimicrobial activity on both isolates, preliminary test by zone of inhibition was done to ascertain their antimicrobial properties and how each extractant impacts this activity. With DMSO as negative control, the extracts were qualitatively assayed for their ability to cause a zone of clearing on each isolate as shown in Figures 1a and 1b. Larger zones were noticed for *S. aureus* than *L. monocytogenes*. This higher size of zone of inhibition observed for *S. aureus* more than those seen for *L. monocytogenes* (p<0.05) could indicate several factors including variable growth patterns of both isolates on Meuller-Hinton agar. 100mg/ml of the extract was sufficient to exhibit antimicrobial activity against both isolates despite the limiting factors faced by this method.

**Figure 1.**
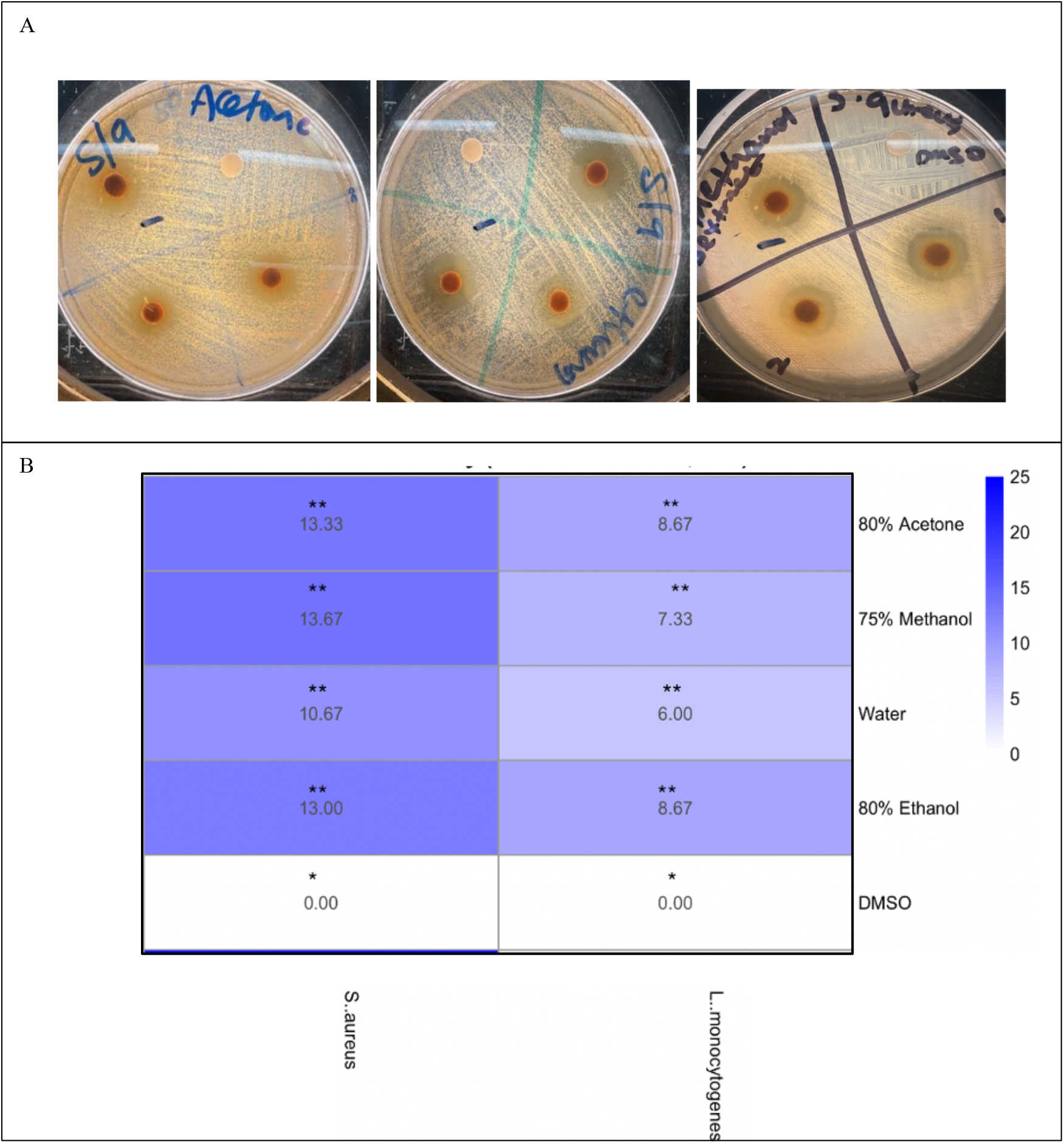
Preliminary test of antibacterial activity of clove extract against S. aureus and L. monocytogenes. (A) Photos of the diameter of the inhibition zone of clove extract of ethanol, methanol, acetone and water on S. aureus and L monocytogenes. (B) The diameter of the inhibition zone by clove extract on both isolates. Number of * indicate significant differences between different extraction methods.

### Quantifying antimicrobial activity via MIC and MBC

The least concentration of the extracts that could inhibit the growth of the isolates was determined via microbroth dilution method following the clinical laboratory standard institute method. The MIC validation was done using the redox ability of resazurin to change from blue to pink color in wells containing viable cells as shown on figure 2a. Acetone and ethanol clove bud extracts showed lower MIC on *S. aureus* which is significantly different from those of water and methanol. Water had the highest MIC among all extraction methods showing a low antimicrobial activity. For *L. monocytogenes*, the acetone and water extract showed higher MIC with ethanol and methanol having the same MIC significantly lower than those of water and acetone (p<0.05) as shown in figure 2b.

**Figure 2.**
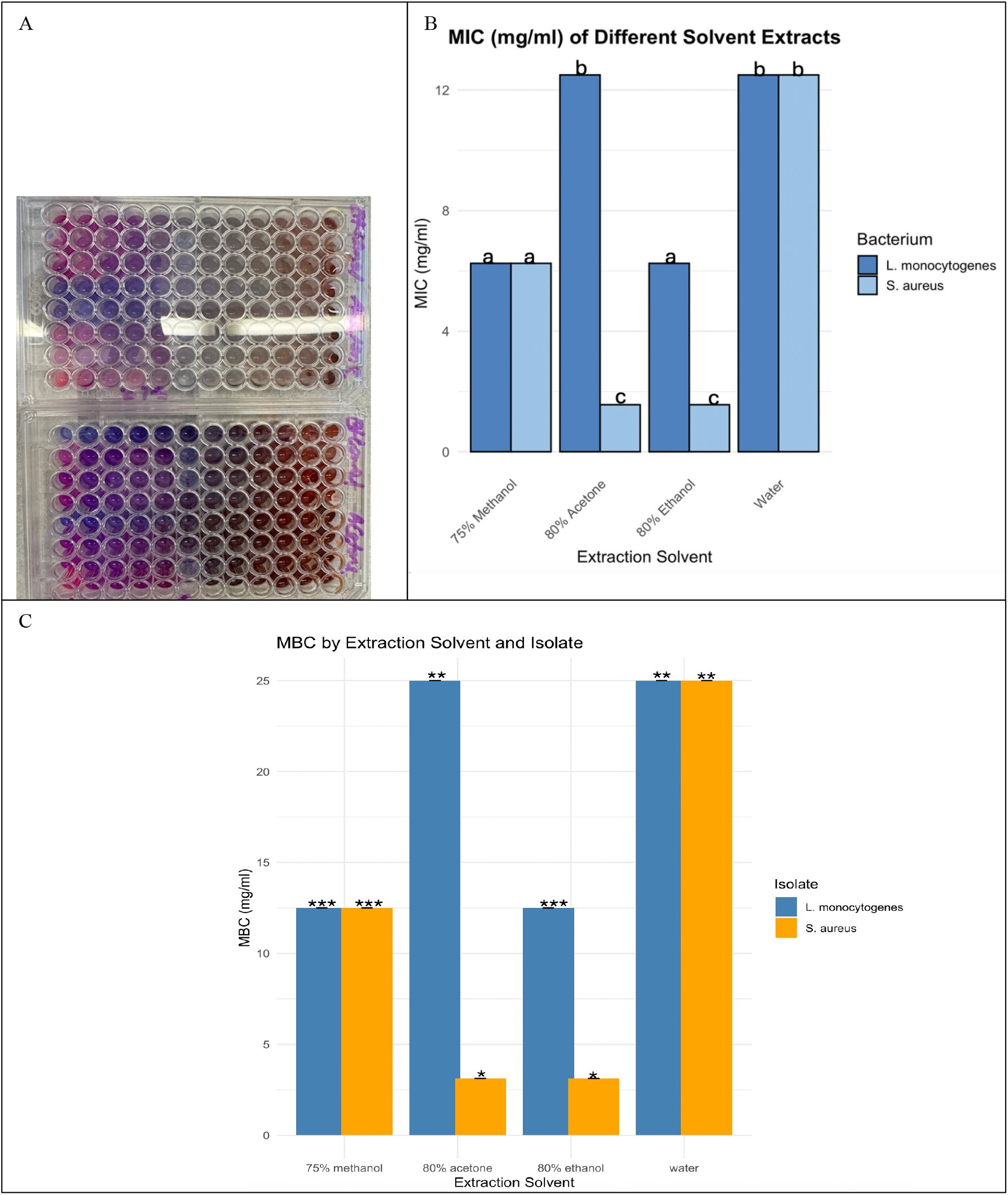
Quantification of antimicrobial activity of crude extract. Figure 2A shows a picture of 96-well plate with wells treated with resazurin. 2B shows the MIC and MBC of different solvents in both isolates. the number of * shows the difference in significant difference. The different letters meat that they are significantly different as p<0.05

To further determine the concentration at which the extract completely kills the viable colonies, the MBC was determined using plates with no visible colonies after incubation. The MBC pattern was the same as MIC for both isolates across all samples but only at a higher concentration of the extract as shown in figure 2c (p<0.05)

### Antimicrobial activity of clove extract is impacted by extraction yield

To determine the correlation between zone of inhibition, MIC and MBC, Pearson correlation analysis was done for both *S. aureus* and *L. monocytogenes*. In both isolates, the correlation between MIC and MBC was 100% as represented by the intensity of the heat map as shown in the figure key and as shown in figure 3A and 3B. The zone of inhibition is negatively correlated with MIC and MBC for both isolates as high zone of inhibition and low MIC indicates a strong activity while a high MIC and low zone of inhibition is an indication of weak activity indicated by the r values. There is a significant difference in the correlation levels for both isolates as stronger correlation was observed for *S. aureus*.

**Figure 3.**
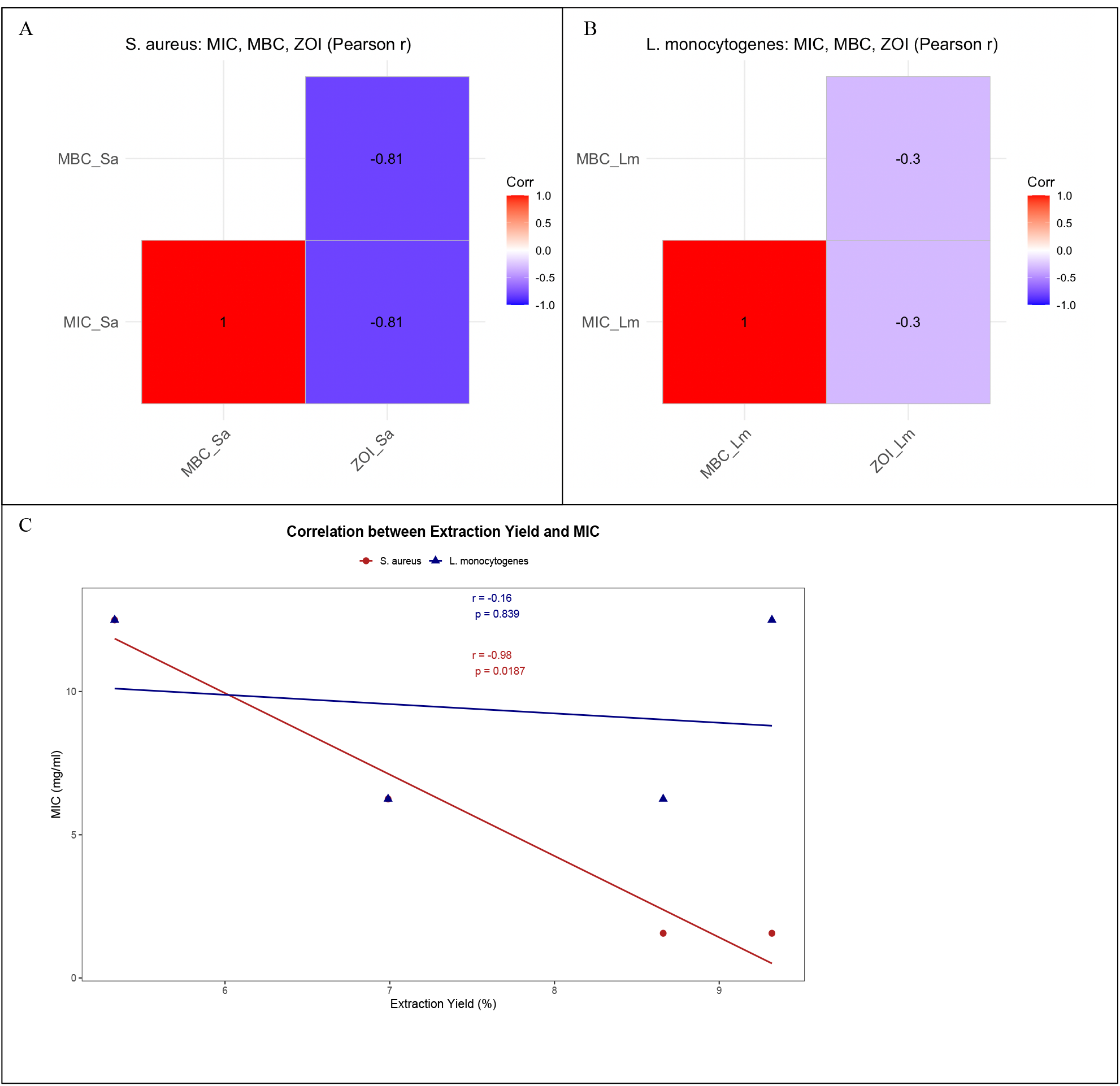
Pearson correlation analysis of antimicrobial activities of the different extraction solvents on *S. aureus* and *L. monocytogenes*. Figure 3A and B shows correlation of MIC, MBC and Zone inhibition in *S. aureus* and *L. monocytogenes*. Figure 3C shows the correlation between extraction yield and MIC.

Pearsson correlation analysis was also carried out to determine how extract yield impacts antimicrobial activity. The r values for antimicrobial effect of the extract on *S. aureus* shows - 0.98 at p<0.05 and -0.16 for *L. monocytogenes* as shown in figure 3C indicating a strong correlation between MIC and extract yield for *S. aureus*.

## Results

## Discussion

The antimicrobial activity of plant extracts has been explored on diverse groups of bacteria including those of foodborne pathogens, but little attention has been paid to the impact of extraction methods on the efficacy of these extracts. (Mytle et al., 2006) assessed the antimicrobial activity of clove oil in inhibiting Listeria monocytogenes H7776 on chicken frankfurters. They discovered that 2% clove oil decreased *L. monocytogenes* within 30 minutes of treatment at very low temperatures. The result from this study shows that solvent polarity has impact on the extraction efficiency and antimicrobial activity of *Syzygium aromaticum* (clove) bud extracts against *Staphylococcus aureus* and *Listeria monocytogenes*. Consistent with published phytochemical extraction principles, 80% acetone produced the highest extraction yield, whereas water yielded the lowest quantity of extractable components. This trend aligns with study by (Kumar et al., 2021)(Liu et al., 2022) which demonstrates that organic solvents with intermediate polarity, particularly aqueous acetone, effectively solubilize phenolic compounds. The major bioactive component of clove is eugenol and its derivative. The low yield obtained from the aqueous extract indicates limited solubility of the hydrophobic and polar components of the extract in water. Mandal et al., 2015). This result is consistent with reports from results reported by (Mandal et al., 2015) displayed by aromatic spices and medicinal botanicals (Mandal et al., 2015) Furthermore, the low significant variation between water and 75% methanol (p > 0.05) may potentially indicate that methanol concentration was not optimally optimized to effectively recover bioactive compounds from clove and sometimes may require acidification corroborating findings from (Hossen et al., 2022) that solvent strength markedly modulates clove phenolic extraction efficiency.

Qualitative screening via agar diffusion revealed clear zones of inhibition for all solvent extracts except the negative control confirming inherent antimicrobial activity of clove extracts. Notably, larger inhibition zones were observed for *S. aureus* relative to *L. monocytogenes* (p < 0.05), indicating greater susceptibility of the Gram-positive cocci to clove extract. Previous study similarly shows increased sensitivity of *S. aureus* to phenolic extracts due to membrane perturbation, enzyme inhibition, and interference with metabolic pathways (Wang et al., 2015) (Chen et al., 2024).The comparatively smaller inhibition zones for *L. monocytogenes* may be attributed to its distinct cell envelope structure, stress response machinery, and biofilm-forming capacity, which collectively confer higher tolerance to phytochemicals (Uddin Mahamud et al., 2024)

Quantitative analysis using broth microdilution confirmed solvent-dependent antimicrobial potency. For *S. aureus*, acetone and ethanol extracts exhibited the lowest minimum inhibitory concentrations (MIC), significantly potent than water and methanol derived extracts. This pattern mirrors solvent-dependent extraction trends, suggesting a direct link between phenolic recovery and bactericidal efficacy. Similar observations have been reported wherein ethanol and acetone extracts of clove demonstrate potent antimicrobial properties attributed to superior solubilization of eugenol and synergistic phenolics(Karaman & Kaplan, 2022). Conversely, water extracts displayed the highest MIC values and thus the weakest antimicrobial efficacy, underscoring that aqueous extraction may fail to isolate key active compounds responsible for antimicrobial action.

Against *L. monocytogenes*, MIC determination showed a slightly different pattern: water and acetone extracts yielded higher MIC values, whereas ethanol and methanol extracts demonstrated comparatively stronger inhibitory activity. This observation suggests that solvent polarity interacts with species-specific sensitivity profiles, possibly reflecting differential permeability of the bacterial cell membrane and varying affinity of phytochemicals to target sites (Adukwu et al., 2012). Minimum bactericidal concentration (MBC) results had same pattern of MIC trends across all solvent treatments, consistent with literature emphasizing that phenolic-rich clove extracts exert both bacteriostatic and bactericidal effects at escalating concentrations (Babu et al., 2011).

Correlation analyses confirmed a robust inverse relationship between inhibition zone size and MIC/MBC values, particularly for *S. aureus* (r = −0.98, p < 0.05). This strong agreement indicates methodological consistency and emphasizes the reliability of the extract’s antimicrobial capacity. In contrast, *L. monocytogenes* demonstrated a weaker correlation (r = −0.16), likely reflecting intrinsic tolerance mechanisms and potential phenotypic variability in stress adaptation. The perfect correlation between MIC and MBC in both organisms further affirms extract potency progression from bacteriostasis to bactericidal activity. Importantly, solvent extraction yield positively correlated with antimicrobial potency against *S. aureus*, reinforcing that quantitative phytoconstituent recovery impacts functional efficacy (Ramesh et al., 2025).

Collectively, these findings strengthen the evidence supporting clove buds as a valuable source of natural antimicrobials with promising utility against Gram-positive pathogens of clinical and food-safety significance. The results corroborate that solvent selection is pivotal in optimizing both extract yield and antimicrobial functionality. Acetone and ethanol emerged as superior solvents for maximizing phenolic yield and antibacterial potency, whereas water demonstrated limited efficacy, reflecting its restricted solubilization of bioactive phytochemicals. The differential susceptibility between the tested organisms underscores the need for species-specific consideration in bio preservative strategies, especially when targeting resilient food-borne pathogens such as *L. monocytogenes*. Future research integrating metabolomic profiling and mechanistic assays is needed to delineate the specific active compounds responsible for observed antimicrobial actions and evaluate their stability in food matrices. Moreover, the conclusion that extraction solvent markedly influences the yield and antimicrobial efficacy of clove bud extracts support calls for the continued exploration of optimized extraction systems and phenolic-rich natural antimicrobials as sustainable alternatives to synthetic preservatives and conventional antimicrobial therapies.

## Acknowledgement

We acknowledge the Department of Microbiology, Federal University of Technology for providing the isolates. We also acknowledge the Anthony Van Leeuwenhoek’s Laboratory for providing all the research supplies and equipment used for this project.

## Author contributions

J. C.H conceived and designed the project. J.C.H and W.B. did the experiment. JCH wrote the manuscript.

